# Evolutionary loss of the ß1-adrenergic receptor in salmonids

**DOI:** 10.1101/2023.01.25.525509

**Authors:** William Joyce

## Abstract

Whole-genome duplications (WGDs) have been at the heart of the diversification of ß-adrenergic receptors (ß-ARs) in vertebrates. Non-teleost jawed vertebrates typically possess three ß-AR genes: *adrb1* (ß1-AR), *adrb2* (ß2-AR), and *adrb3* (ß3-AR), originating from the ancient 2R (two rounds) WGDs. Teleost fishes, owing to the teleost-specific WGD, have five ancestral *adrb* paralogs (*adrb1, adrb2a, adrb2b, adrb3a* and *adrb3b*). Salmonids are particularly intriguing from an evolutionary perspective as they experienced an additional WGD after separating from other teleosts. Moreover, adrenergic regulation in salmonids, especially rainbow trout, has been intensively studied for decades. However, the repertoire of *adrb* genes in salmonids has not been yet characterized. An exhaustive genome survey of diverse salmonids, spanning five genera, complemented by phylogenetic sequence analysis, revealed each species has seven *adrb* paralogs: two *adrb2a*, two *adrb2b*, two *adrb3a* and one *adrb3b*. Surprisingly, salmonids emerge as the first known jawed vertebrate lineage to lack *adrb1. adrb1* is nevertheless highly expressed in the hearts of non-salmonid teleosts, indicating that the wealth of data on adrenergic regulation in salmonids should be generalised to other teleost fishes with caution. It is hypothesised that the loss of *adrb1* could have been viable because of the evolutionary radiation of *adrb2* and *adrb3* genes attributable to the salmonid WGD.

## 1. Introduction

ß-adrenergic receptors (ß-ARs) are G-protein coupled receptors (GPCRs) responsible for transducing the signal of adrenaline and noradrenaline, released as neurotransmitters and hormones, to activate intracellular signalling cascades (Gurevich, 2023; Vasudevan et al., 2011). In most jawed vertebrates, including mammals, there are three ß-ARs; ß1-ARs, ß2-ARs and ß3-ARs, encoded by *adrb1, adrb2* and *adrb3* genes, respectively (Aris-Brosou et al., 2009; Wang et al., 2023; Zavala et al., 2017). The existence of the three ß-ARs, which are differentially expressed in different tissues, allows multifaceted responses to adrenergic stimulation (Gurevich, 2023).

The three main *adrb* paralogs arose from a single gene that multiplied in the ‘2R’ (two rounds) whole-genome duplications (WGDs) in the ancestor of jawed vertebrates (Aris-Brosou et al., 2009; Wang et al., 2023; Zavala et al., 2017). The ancestral fourth *adrb* gene was lost soon after its origin and is not retained in any extant lineages (Zavala et al., 2017). Teleost fishes underwent an additional (‘3R’) WGD after diverging from other ray-finned fishes (Amores et al., 1998; Davesne et al., 2021; Dornburg and Near, 2021; Glasauer and Neuhauss, 2014) which has endowed teleosts with numerous expanded gene families, including *adrb* genes (Zavala et al., 2017). The duplicated copy of *adrb1* is not found in any extant lineages so was apparently lost shortly after the teleost-specific WGD, whereas duplicated *adrb2* (*adrb2a* and *adrb2b*) and *adrb3* (*adrb3a* and *adrb3b*) are retained in some modern lineages (Zavala et al., 2017), such as zebrafish (*Danio rerio*) (Wang et al., 2009).

The salmonid lineage (Salmonidae family: salmon, trout, whitefishes, and their relatives) experienced a fourth (‘4R’) WGD which had an indelible impact on their genomic architecture (Berthelot et al., 2014; Lien et al., 2016; Macqueen and Johnston, 2014). However, the effect of the salmonid-specific WGD on *adrb* gene evolution has not been received dedicated attention. In the most recent and authoritative investigation on *adrb* evolution, no salmonid species were included (Zavala et al., 2017), and whilst earlier phylogenetic analyses included select salmonid *adrb* sequences (Aris-Brosou et al., 2009; Giltrow et al., 2011; Nickerson et al., 2003; Wang et al., 2009), they were not yet able to include a systematic and comprehensive genome-wide survey. This deficit is remarkable given that salmonids, particularly rainbow trout, have for decades served as workhorse model organisms in comparative studies aimed at understanding the mechanisms and importance of adrenergic control in fish (*e*.*g*. Abramochkin et al., 2022; Aho and Vornanen, 2001; Ask et al., 1980; Gamperl et al., 1994; Gesser et al., 1982; Gilbert et al., 2019; Llach et al., 2004; Lortie and Moon, 2003; Miller et al., 2011; Milligan, 1991; Nickerson et al., 2003; Petersen et al., 2013; Sandblom and Axelsson, 2006; Shiels and Farrell, 1997; Vornanen, 1998; Wood and Shelton, 1975). Amongst the most salient findings, whilst ß1-adrenergic receptors are known to be highly expressed in the hearts other vertebrates, especially tetrapods (Abraham et al., 2019; Rohrer et al., 1996), the rainbow trout heart is understood to express virtually exclusively ß2-ARs (Gamperl et al., 1994). In addition, rainbow trout red blood cells were initially believed to express a ß1-AR (Tetens et al., 1988) but later gene expression and cloning experiments identified it to be a ß3-AR (Nickerson et al., 2003). It has been suggested ß1-adrenergic receptors may have a role in immune control in salmonids (*e*.*g*. leucocytes) (Owen et al., 2007) although this largely stems from inferences from other teleost species.

The whole-genome sequencing of various salmonid species (Berthelot et al., 2014; Lien et al., 2016; Mérot et al., n.d.; Smith et al., 2022) has made it possible to now investigate the evolution of their *adrb* genes. The goal of the present study was to establish the *adrb* repertoire of salmonid fishes in the context of the salmonid-specific WGD in order to provide a better foundation for the understanding of adrenergic regulation in fishes.

## 2. Methods

### 2.1 Survey of genomic databases

Protein sequences were first attained from the National Center for Biotechnology Information (NCBI) database. The following species were *a priori* chosen based on their key phylogenetic positions, the availability of well-annotated genomes, and/or their relevance as experimental organisms: one cartilaginous fish, elephant shark (*Callorhinchus milii*); one tetrapod, human (*Homo sapiens*); two non-teleost ray-finned fishes, gray bichir (*Polypterus senegalus*) and spotted gar (*Lepisosteus oculatus*); four non-salmonid teleosts, zebrafish (*Danio rerio*), Atlantic cod (*Gadus morhua*), Japanese medaka (*Oryzias latipes*) and Northern pike (*Esox lucius*); and five salmonids, lake whitefish (*Coregonus clupeaformis*), Atlantic salmon (*Salmo salar*), brown trout (*Salmo trutta*), rainbow trout (*Oncorhynchus mykiss*) and lake trout (*Salvelinus namaycush*). *E. lucius* is a member of the sister lineage (Esociformes) to Salmonidae (Hughes et al., 2018). *C. clupeaformis* belongs to the earliest diverging salmonid lineage relative to the other included species (Hughes et al., 2018; Lien et al., 2016; Ma et al., 2016; Macqueen and Johnston, 2014). ß-AR protein sequences associated with *adrb* genes were found by using basic local alignment search tool (BLAST) (Altschul et al., 1990), *i*.*e*. the nucleotide blast (‘blastn’) and protein blast (‘blastp’) suites (BLAST+ 2.13.0), using default algorithm settings, which were queried using the five zebrafish and three human *adrb*/*ADRB* genes. Searches were performed against the NCBI database nucleotide collection (nr/nt) and non-redundant protein sequences (nr) with results limited to a given species under the ‘organism’ selection. The searches were exhaustive in that, beyond *adrb* genes, they also returned hits for other GPCRs (*i*.*e*. α-adrenergic and serotonin receptors) with lower scores.

To address the possibility that *adrb* genes had not been properly included in the genome assemblies, ‘blastn’ was also performed against unassembled whole-genome sequencing reads in the NCBI Sequence Read Archive for *S. salar* (SRX518053, SRX539891, SRX518057, SRX518056) and *C. clupeaformis* (SRX14636785, SRX14636784). To increase the likelihood of finding *adrb1* matches in salmonids, *E. lucius adrb1* mRNA (XM_010905300.3) was used as query. All returned hits with a close similarity (defined as ‘% identity’ above 70% and a BLAST ‘Total Score’ above 50) were screened by searching for the sequence of the read as query against the NCBI database (‘blastn’ against nucleotide collection nr/nt). Each of the reads yielded a (virtually) identical match against an already identified *adrb* gene for each species (typically 100% identity although occasionally [< 10% reads] with 1 nucleotide mismatch attributable to minor sequencing errors). In both species, the range of hits included matches with the entire set of previously identified *adrb* genes. This leaves it unlikely that any salmonid *adrb* gene, especially an *adrb1* ortholog, was missed in the searches of the NCBI genome assemblies.

The NCBI searches were then cross-checked using the Ensembl genome browser (Cunningham et al., 2022), which presently includes 65 species of ray-finned fishes. Orthologs of human *ADRB1* (ENSG00000043591), *ADRB2* (ENSG00000169252) and *ADRB3* (ENSG00000188778) were retrieved. For the three salmonids represented in both the NCBI search and on Ensembl, *S. salar, S. trutta* and *O. mykiss*, seven *adrb* paralogs per species were identified on Ensembl (four *ADRB2* orthologs and three *ADRB3* orthologs), which corresponded one-to-one with the seven *adrb* genes collected for each species *via* NCBI. Ensembl also provided sequences (also four *ADRB2* orthologs and three *ADRB3* orthologs) for an additional salmonid species, the huchen (*Hucho hucho*), that was not available on the NCBI database. *H. hucho* is sister to the *Oncorhynchus*/*Salvelinus*/*Salmo* clade (Lien et al., 2016; Ma et al., 2016; Macqueen and Johnston, 2014). The Ensembl hits also accounted for all of the *adrb* genes found in the non-salmonid teleosts, zebrafish (*D. rerio*) (*ADRB1*: 1 (gene), *ADRB2*: 2 (genes), *ADRB3*: 2 (genes)), Atlantic cod (*G. morhua*) (*ADRB1*: 1, *ADRB2*: 2, *ADRB3*: 1), Japanese medaka (*O. latipes*) (*ADRB1*: 1, *ADRB2*: 2, *ADRB3*: 1) and Northern pike (*E. lucius*) (*ADRB1*: 1, *ADRB2*: 2, *ADRB3*: 2). Surveys were performed in January 2023.

### 2.2 Phylogenetic analysis

The protein sequences collected through the NCBI and Ensembl searches were compiled to generate an alignment. As outgroups, human dopamine receptors D1 and D5 (*DRD1* and *DRD5*) were used to root the tree, and *adrb*-like sequences from two amphioxus species (*Branchiostoma floridae* and *B. lanceolatum*) and a tunicate (*Ciona intestinalis*) were also included, as based on previous work (Zavala et al., 2017). The final dataset was composed of 77 sequences. The alignment was generated with MAFFT (v7.511) using the L-INS-i strategy (Katoh and Standley, 2013). A maximum likelihood phylogenetic tree was generated with W-IQ-TREE (Trifinopoulos et al., 2016) using the ModelFinder (Kalyaanamoorthy et al., 2017) with default parameters (best-fit model ‘JTT+I+G4’ selected using the Bayesian Information Criterion). To assess node support, ultrafast bootstrap support was calculated using 1000 bootstrap alignments (Hoang et al., 2018). The maximum likelihood tree was exported and visualised in FigTree v1.4.4 (http://tree.bio.ed.ac.uk/software/figtree/).

### 2.3 Conserved synteny analysis

To provide broader insight into the possible absence of *adrb1* in salmonids, conserved synteny diagrams were generated in Genomicus v108.01 using PhyloView (Nguyen et al., 2022). Human *NHLRC2* (which neighbours *adrb1*/*ADRB1* in diverse jawed vertebrates (Wang et al., 2009; Zavala et al., 2017)) was used as the reference gene to provide comparisons with elephant shark, select teleosts (including salmonids) and spotted gar. The salmonids available on Genomicus only included representatives from the Salmoninae sub-family (*S. salar, S. trutta, O. mykiss, Oncorhynchus kisutch*. Note that *Hucho hucho*, another Salmoninae member (Macqueen and Johnston, 2014), was also available but it was excluded from the final figure as the available genomic region only included a small chromosome fragment which was nevertheless consistent with the salmonid species that had greater coverage). Therefore, the NCBI gene page for lake whitefish (*C. clupeaformis*) (www.ncbi.nlm.nih.gov/gene/121547156/), representative of an earlier-diverging salmonid lineage (Lien et al., 2016; Macqueen and Johnston, 2014), was additionally consulted. The ‘genomic context’ data confirmed a similar arrangement of genes neighbouring *nhlrc2* as the Salmoninae species available on Genomicus, which is therefore inferred as ancestral to the Salmonidae family.

### 2.4 Survey of a transcriptomic database

To further investigate if *adrb1* genes can be found in salmonids the PhyloFish transcriptomic database (Pasquier et al., 2016) was used. This includes assembled transcriptomes for 23 species of phylogenetically diverse ray-finned fish including 6 salmonids: bowfin (*Amia calva*), spotted gar (*L. oculatus*), European eel (*Anguilla anguilla*), butterflyfish (*Pantodon buchholzi*), arowana (*Osteoglossum bicirrhosum*), elephantnose fish (*Gnathonemus petersi*), allis shad (*Alosa alosa*), zebrafish (*D. rerio*) panga (*Pangasianodon hypophthalmus*), black ghost knifefish (*Apteronotus albifrons*), Mexican tetra (*Astyanax mexicanus*), Northern pike (*E. lucius*), Eastern mudminnow (*Umbra pygmae*), grayling (*Thymallus thymallus*), (European/)common whitefish (*Coregonus lavaretus*), (American/)lake whitefish (*C. clupeaformis*), brown trout (*S. trutta*), rainbow trout (*O. mykiss*), brook trout (*Salvelinus fontinalis*), ayu (*Plecoglossus altivelis*), Atlantic cod (*G. morhua*), Japanese medaka (*O. latipes*) and European perch (*Perca fluviatilis*). The database was first queried with the zebrafish ß1-AR protein sequence (NP_001122161.1) using the ‘tblastn’ function. Transcripts were recorded, starting with those with the highest similarity to NP_001122161.1, until the search began to identify *adrb2* and *adrb3* paralogs in species that had already retuned a hit for *adrb1*. The search was also repeated with pike ß1-AR protein sequence (XP_010903602.1) as query, which identified an identical set of *adrb1* transcripts. To check that the identified transcripts corresponded to *adrb1* sequences, the nucleotide sequences were translated to protein sequences using the Expasy translate tool (web.expasy.org/translate/) and the phylogenetic analysis (as described above) was repeated with their inclusion. This process verified all of the sequences retrieved from PhyloFish were phylogenetically clustered (ultrafast bootstrap support of Actinopterygii *adrb1*: 99%) with other vertebrate *adrb1* sequences so are presumably orthologs.

The PhyloFish database was also employed to estimate *adrb* expression levels of the heart in all of the non-salmonids used in the phylogenetic analysis that were available (*i*.*e*. spotted gar (*L. oculatus*), zebrafish (*D. rerio*), Atlantic cod (*G. morhua*), Japanese medaka (*O. latipes*) and Northern pike (*E. lucius*)) as well as all six available salmonids: grayling (*T. thymallus*), European whitefish (*C. lavaretus*), lake whitefish (*C. clupeaformis*), brown trout (*S. trutta*), rainbow trout (*O. mykiss*) and brook trout (*S. fontinalis*). PhyloFish was searched, as described above, using the ‘tblastn’ function for with the known *adrb* sequences of each paralog. Transcriptome count files were downloaded from PhyloFish and abundance was calculated in transcripts per million (TPM).

## 3. Results and Discussion

### 3.1. Genomic survey and phylogenetic classifications of *adrb* subgroups

To investigate the effects of the salmonid specific-WGD (Berthelot et al., 2014; Lien et al., 2016; Macqueen and Johnston, 2014) on ß-AR evolution, the genomes of six salmonids representing five genera (lake whitefish (*Coregonus clupeaformis*), huchen (*Hucho hucho*), Atlantic salmon (*Salmo salar*), brown trout (*Salmo trutta*), lake trout (*Salvelinus namaycush*), and rainbow trout (*Oncorhynchus mykiss*)) were exhaustively searched for *adrb* genes using NCBI and Ensembl databases. In each of these species seven *adrb* genes were identified. On the NCBI database, this included sequences that have been provisionally attributed as ß1, ß2 and ß3-AR(-like). Phylogenetic analysis and conserved synteny were used to clarify the classifications of the salmonid *adrb* genes by providing comparisons with previously characterized species.

Providing a scaffold for the integration of salmonid *adrb* sequences, the non-teleost jawed vertebrates included in the study (elephant shark, human, bichir and spotted gar) consistently had three readily identifiable *adrb* genes (*adrb1, adrb2, adrb3*), as shown previously for the vast majority of gnathostome lineages (Aris-Brosou et al., 2009; Zavala et al., 2017). As expected due to the teleost-specific whole-genome duplication (Zavala et al., 2017), teleosts demonstrated an increased *adrb* complement. Of the species included here, zebrafish and pike retain all five ancestral *adrb* genes (*adrb1, adrb2a, adrb2b, adrb3a* and *adrb3b*) whereas medaka and cod lost *adrb3b*, resulting in a final repertoire of four *adrb* genes (*adrb1, adrb2a, adrb2b* and *adrb3a*).

Based on phylogenetic clustering with annotated zebrafish genes, each salmonid species exhibited two *adrb2a* paralogs, two *adrb2b* paralogs, two *adrb3a* paralogs, and one *adrb3b* paralog (Fig. 1), indicating versions of these seven genes were most likely shared in their last common ancestor. Because pike (*Esox lucius*, Esocidae family), a member of the sister lineage to Salmonidae that diverged prior to the salmonid-specific WGD (Berthelot et al., 2014; Hughes et al., 2018; Lien et al., 2016; Macqueen and Johnston, 2014), only has a single *adrb2a, adrb2b* and *adrb3a* paralog, each acting as sister to a clade of duplicate salmonid paralogs, the two copies of each in salmonids can be ascribed to their 4R WGD. Furthermore, in Atlantic salmon (*S. salar*), the genomic locations of the duplicates of *adrb2a* (chromosomes 5 and 9), *adrb2b* (chromosomes 4 and 13) and *adrb3a* (chromosomes 20 and 24) correspond seamlessly with the homeologous genomic syntenic blocks mapped and attributed to the salmonid-specific WGD by Lien et al. (2016).

**Figure 1.**
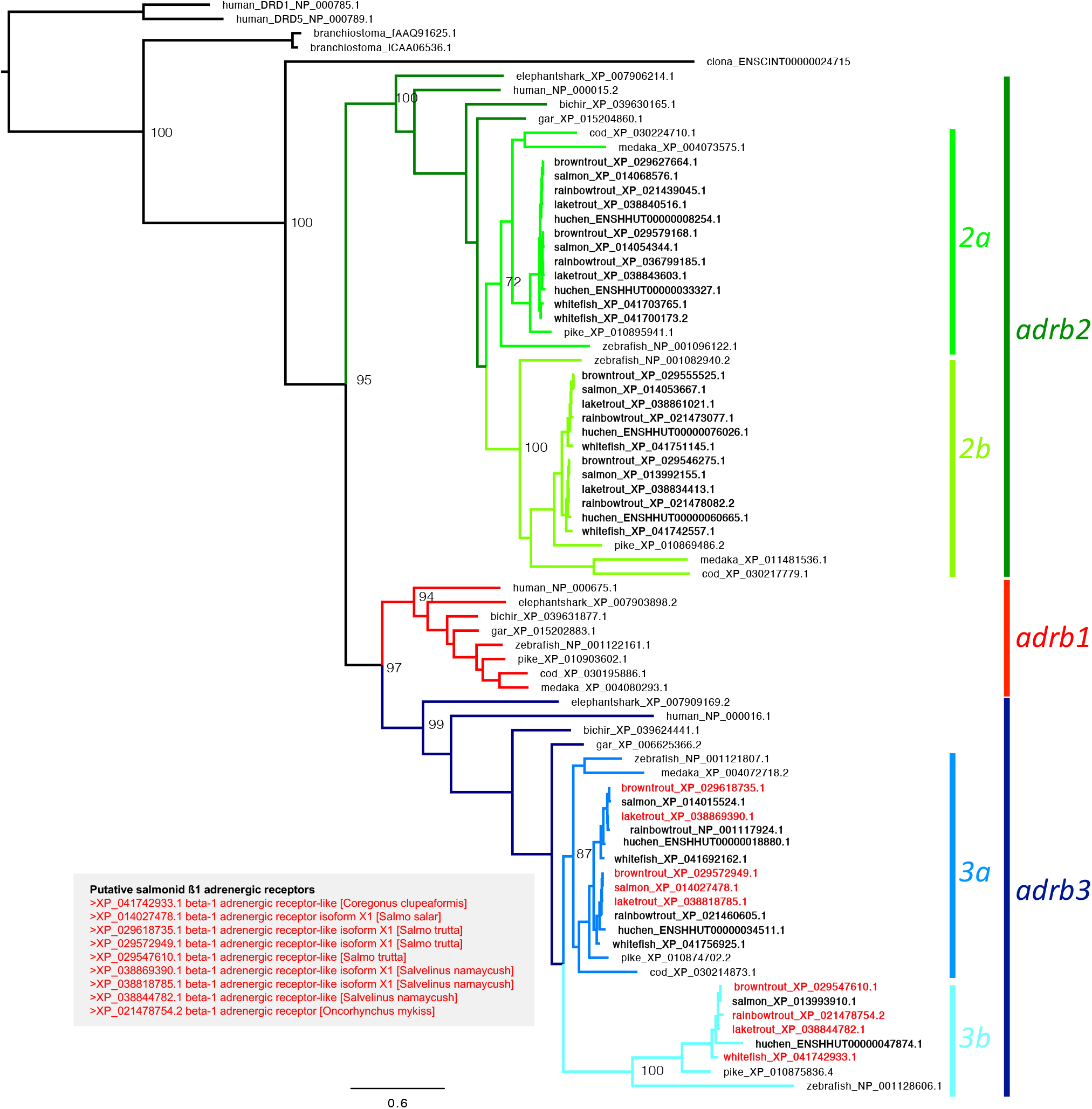
Maximum likelihood phylogenetic tree of vertebrate protein sequences associated with *adrb* genes, with a focus on salmonid fishes. The tree was generated using W-IQ-TREE. Attribution to *adrb* classes *(adrb1, adrb2*(*a*/*b*), *adrb3*(*a/b*)) was performed based on presence of human and zebrafish sequences in a monophyletic cluster. Salmonids are indicated with bolded labels. Labels in red text and inset box shows proteins currently identified as ß1-adrenergic receptor(-like) on the NCBI database (implying status as *adrb1* genes), all of which were instead recovered as *adrb3*. Species abbreviations are as follows: ‘human’ (*Homo sapiens*); ‘elephant shark’ (*Callorhinchus milii*); (gray) ‘bichir’ (*Polypterus senegalus*) and (spotted) ‘gar’ (*Lepisosteus oculatus*); ‘zebrafish’ (*Danio rerio*), (Atlantic) ‘cod’ (*Gadus morhua*), (Japanese) ‘medaka’ (*Oryzias latipes*) and (Northern) ‘pike’ (*Esox lucius*); (lake) ‘whitefish’ (*Coregonus clupeaformis*), (Atlantic) ‘salmon’ (*Salmo salar*), ‘brown trout’ (*Salmo trutta*), ‘rainbow trout’ (*Oncorhynchus mykiss*) and ‘lake trout’ (*Salvelinus namaycush*); The tree is rooted with human dopamine receptors D1 and D5 (DRD1 and DRD5) and other outgroups included two amphioxus species (*Branchiostoma floridae* and *B. lanceolatum*) and a tunicate (*Ciona intestinalis*) (Zavala et al., 2017). Values next to nodes show ultrafast bootstrap support (%).

It is noteworthy that both duplicated *adrb2a, adrb2b* and *adrb3a* paralogs were retained in each of the descendent salmonid lineages, given that loss of redundant genes is commonplace after WGD, although retention of duplicated paralogs is relatively typical in salmonids (Berthelot et al., 2014). The rainbow trout *adrb3a* and *adrb3b* genes as cloned and named by Nickerson et al. (Nickerson et al., 2003) (XP_021460605.1 and NP_001117924.1, respectively in Fig. 1) were both recovered as part of the *adrb3a* family (according to standard nomenclature generated from zebrafish genome annotation), as has previously been suggested (Wang et al., 2009), confirming that they originated in the salmon-specific WGD. This reveals an ambiguity with regard to the *adrb3b* nomenclature as deriving from the teleost-specific WGD or the salmonid-specific WGD.

Most surprisingly, no *adrb1* paralog could be identified in any of the salmonid genomes. Each of the salmonid proteins currently described as being ß1-AR(-like) on the NCBI database were invariably recovered as an *adrb3a* or *adrb3b* paralog (Fig. 1). This annotation problem was not unique to salmonids; *adrb3* genes in cod (XP_030214873.1), gar (XP_006625366.2) and elephant shark (XP_007909169.2) are also currently mislabelled as ß1-AR(-like), but these species also have a bona fide *adrb1* paralog elsewhere in their genomes. Evidently, in salmonids, the additional suite of *adrb3* genes generated by their 4R WGD has previously obscured the apparent loss of the true *adrb1*. The lack of *adrb1* in salmonids is consistent with previous phylogenetic studies that have incorporated some salmonid *adrb* sequences but failed to include *adrb1* (Aris-Brosou et al., 2009; Giltrow et al., 2011; Wang et al., 2009) whilst the gene is otherwise broadly found in teleosts (Zavala et al., 2017). Because none of the earlier studies were able to cover all genome-wide *adrb* paralogs in salmonids (each earlier study also lacks the complete set of *adrb2* and *adrb3* paralogs), the realisation that *adrb1* has been lost has not previously been possible.

### 3.2. Conserved synteny

The hypothesis that *adrb1* has been lost in salmonids is additionally supported by analysis of conserved synteny, *i*.*e* preserved co-localisation of genes on a chromosome. In other gnathostome vertebrates, including other teleosts (Fig. 2), *adrb1* is found alongside *tdrd1, ccdc186, nhlrc2* and *dclre1a* genes (Zavala et al., 2017). However, whilst these four genes are still found in proximity in salmonids, there is no neighbouring *adrb* gene present (Fig. 2). Mechanistically, the loss of *adrb1* appears likely to have coincided with the inversion of *tdrd1* and *ccdc186* relative to *nhlrc2* and *dclre1a* (Fig. 2).

**Figure 2.**
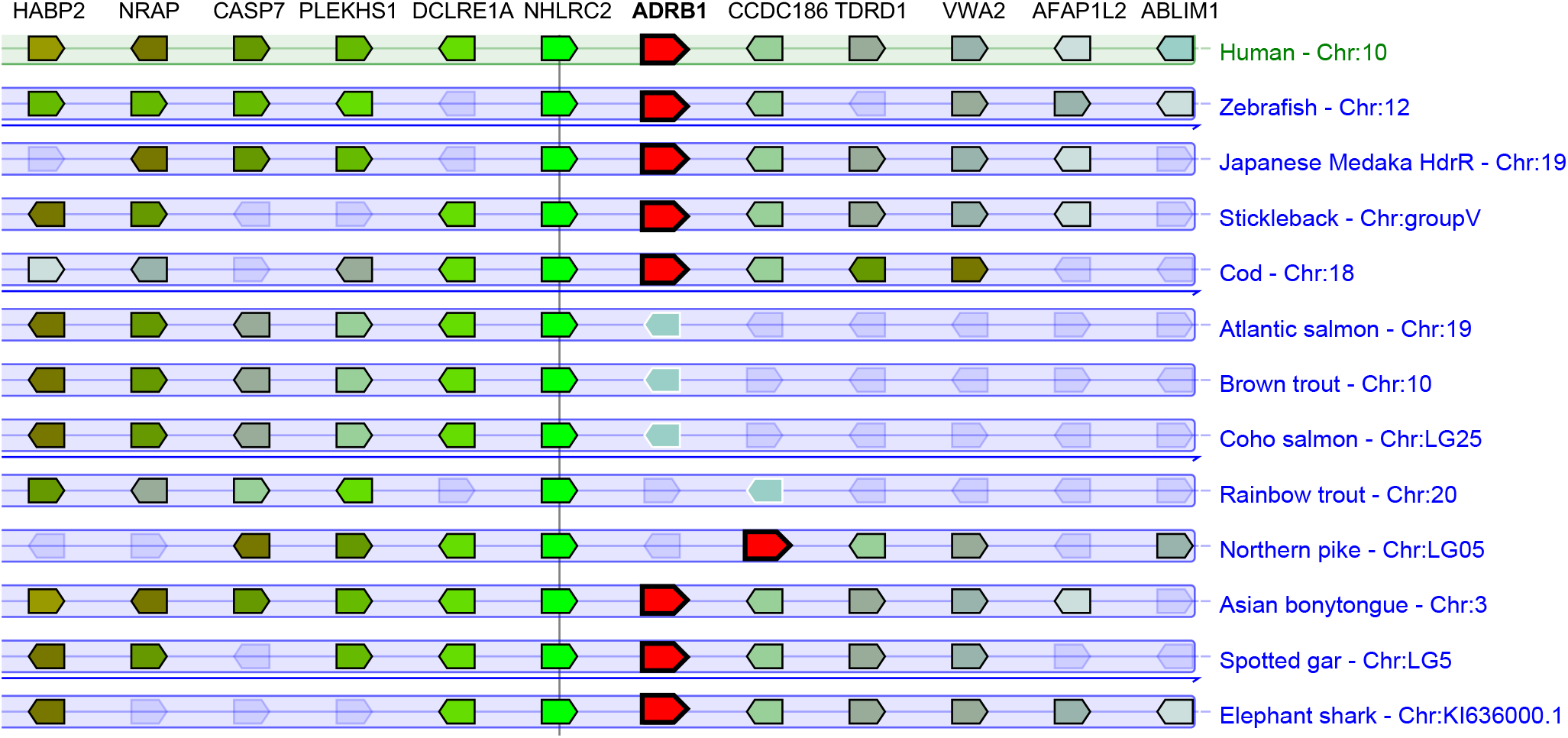
Conserved synteny of the genomic region normally harbouring *ADRB1* (*adrb1* in fishes). Each horizontal pentagon represents a gene, wherein orthologs are shown by a consistent colour. The figure was generated using Genomicus (Nguyen et al., 2022).

### 3.3. Transcriptomic survey and cardiac gene expression

Genomic searches for a given gene are vulnerable to insufficient reference genome sequencing quality, but transcriptomics can provide an alternative avenue to search for otherwise missing genes (*e*.*g*. Yin et al., 2019). The PhyloFish database, including transcriptomes for 23 phylogenetically diverse species of bony fish, including 6 salmonids, was queried with zebrafish the *adrb1* sequence. Providing further evidence for the loss of *adrb1* in salmonids, *adrb1* transcripts were identified in 15/17 non-salmonid bony fish species (all except Eastern mudminnow (*U. pygmaea*) and butterflyfish (*P. buchholzi*)) but 0/6 salmonids (grayling (*T. thymallus*), European whitefish (*C. lavaretus*), American whitefish (*C. clupeaformis*), brown trout (*S. trutta*), rainbow trout (*O. mykiss*), brook trout (*S. fontinalis*)). Due to an absence of genomic data, it is not yet possible to verify if the lack of *adrb1* transcripts in *U. pygmaea* and *P. buchholzi* represents true genomic loss of the encoding gene or simply exhibited too low expression in this transcriptome, with the latter remaining a distinct possibility.

To provide context on the implications of the loss of *adrb1* in salmonids, *adrb* expression in the heart, an organ where the expression of different ß-ARs is of particular focus (Gamperl et al., 1994; Gurevich, 2023; Shiels, 2017), was quantitatively compared across several species of ray-finned fishes (Fig. 3). Consistent with previous cardiac radioligand binding data in rainbow trout showing a predominance of ß2-ARs (Gamperl et al., 1994), *adrb2a* transcripts were found in great abundance in the hearts of diverse salmonids. This was generally complemented by the expression of *adrb3a*, which has previously been documented to be relatively high in rainbow trout heart (Nickerson et al., 2003). However, most of the other fish, including pike, exhibited high transcript counts of *adrb1*, albeit in some cases (*i*.*e*. zebrafish and medaka) in combination with a highly expressed *adrb2* gene (Fig. 3). Abundant expression of *adrb1* has also previously been reported in the hearts of non-salmonid teleosts using quantitative PCR (Giltrow et al., 2011; Steele et al., 2011) and is a characteristic shared with the hearts of other vertebrates, including mammals (Myagmar et al., 2017). Clearly, one should be cautious to infer protein levels from transcript counts, yet this analysis convincingly demonstrates that *adrb1* is expressed in the hearts of non-salmonid fishes. As such, it would be wise to apply caution when extrapolating studies of adrenergic regulation in salmonids to other teleost fish species.

**Figure 3.**
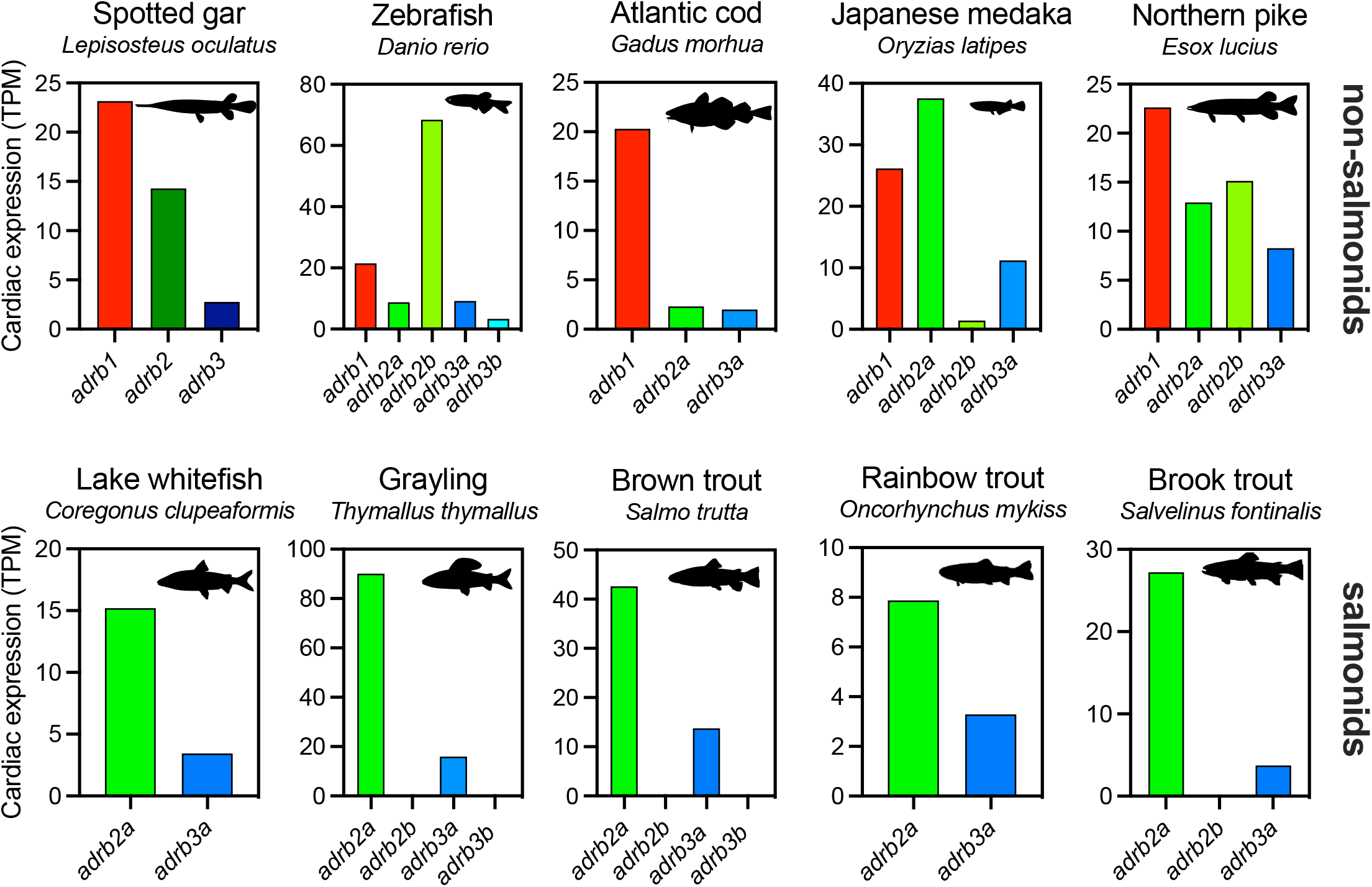
Cardiac *adrb* gene expression. Transcriptomes were downloaded from the PhyloFish database (Pasquier et al., 2016). Counts are expressed as transcripts per million (TPM). In non-salmonids, cardiac *adrb1* expression is typically highly expressed, whereas after the loss of *adrb1* salmonids predominantly express an *adrb2a* paralog in the heart. Fish silhouettes courtesy of http://phylopic.org/ or custom generated from images in the public domain.

### 3.4. Evolutionary implications and conclusions

Through a variety of complementary approaches, this study characterizes salmonids as the first known jawed vertebrate lineage to seemingly lack an *adrb1* gene. In mice, deletion of *Adrb1* results in high prenatal mortality and renders the heart insensitive to ß-adrenergic stimulation (Rohrer et al., 1996). In zebrafish, by contrast, *adrb1* knockout has no obvious adverse effects on viability and fertility, at least when raised in optimized aquaculture conditions (Joyce et al., 2022), although it is not known if the knockout fish would survive under natural conditions. Routine heart rate is suppressed in *adrb1* knockout larval zebrafish, although they are still able to respond to ß-adrenergic stimulation with tachycardia, presumably mediated by other ß-ARs (Joyce et al., 2022).

It is not easily discernible if *adrb1* was lost before or after the salmonid-specific WGD, with both events occurring after the separation of Salmonidae and Esociformes (∼130 million years ago) but pre-dating the divergence of Salmoninae and Coregoninae (∼55 million years ago) (Kumar et al., 2022; Lien et al., 2016). Indeed, the most parsimonious explanation would be loss before the WGD, which would require loss of one, not two, *adrb1* genes. However, it appears plausible that loss of *adrb1* might not have been sustainable without the expansion of *adrb2* and *adrb3* repertoires. By analogy, knockout of *adrb1* in zebrafish leads to increased cardiac expression of *adrb2b* and *adrb3b* genes (Joyce et al., 2022). As such, this study contributes to the growing appreciation of the redundancy between *adrb* paralogs, particularly in teleosts. More generally, the loss of *adrb1* in salmonids provides a compelling example that even when duplicate new paralogs deriving from a recent WGD are retained (Berthelot et al., 2014), the redundancy generated could enable losses of other, more distantly related paralogs in a gene family.

## Acknowledgments

I am grateful to Steve F. Perry, Holly A. Shiels, Daniel M. Ripley and Laura Cadiz for insightful discussions and comments on the manuscript. I also thank Cédric Cabau for advice regarding analysis of the PhyloFish transcriptome database.

